# To group or not to group: Effect of prolonged exposure to predation and resource availability on the shoaling behaviour of zebrafish (*Danio rerio)*

**DOI:** 10.1101/2020.07.02.183897

**Authors:** Nawaf Abdul Majeed, Vivek Philip Cyriac, Ullasa Kodandaramaiah

## Abstract

Individuals of many species live in groups to obtain anti-predatory advantages, foraging benefits and for social reasons. Living in large groups can reduce predation, but as group size increases, competition for resources also increases. The trade-off between the advantages of group living for an individual and competition caused by it can determine group dynamics, and this trade-off can vary with environmental conditions. Shoaling behaviour, the tendency of fish to form groups, is shown to be affected by factors such as resource availability, presence of predators and conspecifics. Although studies indicate that both predation and starvation pressure in an environment can determine whether fish choose to shoal or not, whether prolonged exposure to such conditions influences shoaling behaviour remains little explored. Here, we test how predation pressure and resource availability may interactively shape the shoaling behaviour of zebrafish (*Danio rerio*) when exposed to combinations of these pressures over a two-week period. We find that shoal size increases with predation and decreases with starvation, and that greater predation pressure increases shoaling tendencies even under reduced food availability. Overall, we show that prolonged exposure to varying predation pressure and resource availability can together influence shoaling tendencies of fish even when such pressures are relaxed.

**Significant statement:** In group living species, group structure and dynamics depend on various intrinsic factors and environmental stressors. Shoaling behaviour in fish, where individuals aggregate to form groups, is shown to be altered with environmental factors such as predation and resource availability. Although studies have examined the effects of these cues on shoaling behaviour, the ecological circumstances experienced by fish could also influence shoaling tendencies. We here show that shoaling behaviour is also shaped by previous experience of fish to predation and food resource availability. We check how shoaling behaviour varies with differences in predation pressure and resource availability after prolonged exposure to these conditions by measuring the shoal size and shoal cohesion in zebrafish. This study illuminates how shoaling tendencies of individuals shaped by the environmental conditions persist even when these environmental pressures are removed.

## INTRODUCTION

Living in groups is an important social behaviour exhibited by many species across various environmental conditions (Krause and Ruxton 2002). Animals form social groups in order to gain individual fitness benefits (Pöysä 1992; Pitcher 1998; Tojo et al. 2005; Mikheev 2009; Vanthournout et al. 2016). Group living can facilitate information transfer among individuals (Krause and Ruxton 2002) and enhance social learning (Helfman and Schultz 1984; Brown and Laland 2003), thus improving an individual’s ability to locate and utilise food sources (Krebs et al. 1972; Pitcher et al. 1982; Magurran and Pitcher 1983; Hintz and Lonzarich 2018). Group living can also have anti-predatory advantages (Alexander 1974; Treherne and Foster 1981; Uetz et al. 2002; Miller and Gerlai 2011; Ioannou et al. 2017). For instance, forming large groups can reduce the overall predation risk for each individual in the group through the predator dilution effect (Hamilton 1971; Foster and Treherne 1981; Treherne and Foster 1982; Ioannou et al. 2012; Olson et al. 2015) or by enhancing predator detection by prey because many eyes are more likely to detect and warn other members of approaching predators (Pitcher and Parrish 1993; Lima 1995; Roberts 1996). Living in larger groups can also enable individuals to use a greater proportion of their time foraging than on vigilance (many eyes hypothesis) (Pulliam 1973; Magurran and Pitcher 1983; Pitcher and Parrish 1993; Li et al. 2009; Olson et al. 2015). Individuals in a group can also benefit from the confusion effect that is created when individuals move rapidly in random directions (Caspers 1983; Krakauer 1995; Tosh et al. 2006; Jeschke and Tollrian 2007; Ioannou et al. 2008; Olson et al. 2013). Additionally, increased aggregation when the prey population size is constant can reduce the number of groups and the encounter rates between predators and prey (Ioannou et al. 2012).

Although group living provides several advantages, it can also generate competition for food, mates and shelter (Pitcher and Parrish 1993; Pitcher 1998). Grouping induced competition for food increases with group size (Street et al. 1984; Hoare et al. 2004; Stenberg and Persson 2005) and can be expensive when resources are scarce (Krause 1993; Krause and Ruxton 2002). For instance, it was shown that food scarcity decreases the optimal group size in primates through increased intra-group competition (Pride 2005). However, how the benefits of reduced predation and the costs of competition together mediate group dynamics and grouping tendency require further investigation.

Most fish species aggregate together to form cohesive social groups called shoals at some stage of their life history (Shaw 1978; Pitcher and Parrish 1993; Ryer and Olla 1995). Studies have shown that shoaling behaviour in fish is advantageous in that it can reduce predation by improving predator detection (Magurran et al. 1985; Godin et al. 1988) or via the dilution effect (Foster and Treherne 1981) or by confusing predators (the confusion effect) by moving rapidly in random directions (termed flash expansion) (Landeau and Terborgh 1986). Shoaling can also facilitate foraging through better detection of food patches (Pitcher et al. 1982; Ranta and Juvonen 1993; Ranta and Kaitala 2006), by allowing individual fish to allocate more time for foraging (Magurran and Pitcher 1983) or through improved social learning (Day et al. 2001). However, a larger shoal size also increases competition for resources (Alexander 1974; Miller and Gerlai 2011). Thus, environmental factors such as the presence of predators and the availability and distribution of resources could have conflicting effects on shoaling tendencies, and influence the properties of a shoal (Hoare et al. 2004). Since natural environments are generally dynamic, individual fish would have to make decisions on whether to form large or small shoals depending on the current ecological circumstances (Pitcher 1998). For instance, shoal size and compactness should increase in the presence of high predation pressure (Magurran and Seghers 1991; Huizinga et al. 2009) but should decrease with increased starvation pressure under low food availability (Miller and Gerlai 2007). Such a trade-off between resource availability and predation pressure is a key factor determining the size and compactness of a shoal (Miller and Gerlai 2011).

Several studies have explored how predation and resource availability can independently influence shoaling in fish (Keenleyside 1955; Magurran 1990; Hager and Helfman 1991; Ashley et al. 1993). A few studies have also looked at how predation pressure and resource availability together influence shoal dynamics (Hoare et al. 2004; Miller and Gerlai 2007). Few studies have explored whether the shoaling tendency is also influenced by the assessment of the risk of predation or starvation in a particular environment based on previous experience. Here, using zebrafish (*Danio rerio*) as a model, we tested how prolonged exposure to predation and starvation pressure together interact with each other to shape shoaling behaviour. We show that prolonged exposure to predation and starvation can influence shoaling behaviour in zebrafish even when such pressures are relaxed. We also show that shoal size increases with predation and decreases with starvation and that predation pressure increases shoaling tendencies even under reduced food availability.

## MATERIALS AND METHODS

### Experimental setup

We designed a 2 × 2 factorial experiment with combinations of varying levels of predation and starvation pressure. Zebrafish were procured from local aquarium stores. Fish were maintained in a 30 × 60 × 30 cm storing aquarium for 10 days before the experiment and fed with pellet fish food (Aquafood™, Chennai, India) twice every day. For the experiment, we used four aquaria of 60 × 50 × 15 cm dimensions for the four treatments. The four treatments were (1) high predation and high food availability (HP-HF) (2) high predation and low food availability (HP-LF) (3) low predation and high food availability (LP-HF) and (4) low predation and low food availability (LP-LF). Each experimental aquarium was covered by a black plastic sheet on the outer walls to minimise external influence. Shoaling behaviour of fish was recorded using video cameras placed above each experimental aquarium, 150 cm from the water surface. Following Miller and Gerlai (2007) we restricted the height of the water column in the experimental aquaria to 10 cm so that shoaling could be measured in a two-dimensional space.

### Experimental protocol

After acclimatizing the fish in the storing aquarium for 10 days, we randomly transferred 20 fish into each of the four experimental aquaria and randomly assigned each aquarium to a specific treatment. The fish were left for a week in their respective experimental aquarium under normal conditions before subjecting them to their respective treatments. Each treatment differed in the combination of high and low predation and food availability. The fish were subjected to their respective conditions for two weeks, after which we video-recorded their shoaling behaviour without the presence of starvation or predation pressures.

The fish were provided food and exposed to predation twice per day, once in the morning (10.30–11.30 hours) and once in the evening (16.00–17.00 hours). We simulated high and low food availability by adding 20 and 10 crushed fish food pellets respectively to the respective aquaria. Predation was simulated by spraying 10 ml of freshly prepared zebrafish skin extracts on the water column in each aquarium. We prepared two concentrations of skin extract to simulate high and low predator pressure. High concentration skin extract was made by crushing and mixing the dissected skin of a euthanized zebrafish with 300 ml water, as done in other studies (Von Frisch 1942; Hoare et al. 2004; Speedie and Gerlai 2008). For the low concentration skin extract, we mixed the crushed fish in 3000 ml of water. Skin extracts were kept refrigerated and used within 24 hours of preparation.

After subjecting the aquaria to the respective food and predation conditions for two weeks, we measured the shoaling behaviour for three days by video recording fish activity for 30 minutes, during which time we did not simulate predation or provide food to any of the treatments. The video recordings were carried out twice per day, one session in the morning (10.30–11.30 hours) and one in the evening (16.00–17.00 hours). To quantify the shoal size, we converted each video recording session into image frames and selected 10 random frames from each session for further analyses. We used ImageJ v1.52 (Schneider et al. 2012) to measure the distance between fish and quantify shoal size. We considered two fish to be part of the same shoal if they were within four body lengths (4BL) from each other (<12 cm between the heads of two zebrafish). The 4BL threshold is a standard measure to determine the shoal size in fish (Pitcher 1983; Pitcher and Parrish 1993; Hoare et al. 2004; Miller and Gerlai 2011) and is based on the elective group size concept where individuals within 4BL of each other are assumed to be able to communicate with each other (Pitcher 1983; Pitcher and Parrish 1993). The bottom surface of each experimental aquaria was marked on all sides with short lines at every 5cm interval using a permanent marker prior to the start of the experiment in order to allow calibration of the distances in ImageJ. To measure shoal size, the distance between fish were manually calculated using ImageJ for all the image frames. To measure shoal cohesion, we manually extracted the spatial position of each fish as (x, y) coordinates (with the corner of the aquarium as the point x=0, y=0; the corner was chosen randomly and we are calculating the differences in the spatial points between fish so the corner will not have any effect on the values) from the 10 randomly selected image frames from each video recording session. We then calculated the distance from each fish to its nearest neighbour using the *spatstat* package (Baddeley and Turner 2005) in R v. 3.5.2 (R Core Team 2016). We conducted the experiment twice following the same protocol, the first set in Feb 2019 and the second in March 2019. Aquaria were reassigned randomly to the treatments in March 2019.

### Statistical analysis

All analyses were carried out in R v. 3.3.2 (R Core Team 2016). To check if shoal size differed across treatments, we calculated the mean shoal size for each of the 10 randomly selected frames in each session from the frequency of fish present in different shoal sizes. Since the distribution of mean shoal size within each treatment and all four treatments combined was non-normal, we used a non-parametric Kruskal-Wallis test to check for a difference among the four treatments. We also used a pairwise Mann-Whitney U test to check for differences between treatments. We grouped different shoal sizes into four categories ranging from 1–5, 6–10, 11–15 and 16–20. Since the count data for fish in the four categories were zero-inflated, we used zero-inflated Poisson regression models using the R package pscl v. 1.5.2 (Jackman 2006) to test the effect of treatment and shoal size category. We calculated the estimated marginal means using the R package *emmeans* v. 1.3.3 (Lenth 2019) to obtain pairwise comparisons between the frequency of fish present in different shoal size categories within and across treatments. We also checked for differences in shoal cohesion across treatments by calculating the nearest neighbour distance (NND) for each fish. We log-transformed the NND to make it normally distributed and used linear mixed effect models (LMM) to check the differences between treatments using the R package *lme4* v 1.1-21 (Bates et al. 2015). We evaluated the fit of two models, one where NND was predicted by treatment and another model where NND was independent of treatment, using Δ Akaike Information Criterion (ΔAIC). We considered ΔAIC between 2 and 10 as moderate support and a value >10 as strong support (Burnham and Anderson 2002). We used the Tukey post-hoc test for pairwise comparisons between the NND values across treatments.

## RESULTS

Mean shoal size varied significantly between all treatment pairs (Fig 1) (Kruskal-Wallis: W=117.69, df= 3, p= <0.001,). The mean shoal size was significantly higher in the HP-HF treatment compared to all other treatments (Mann-Whitney U test: HP-HF vs HP-LF: p = 4.4e-11; HP-HF vs LP-HF: p = <2e-16; HP-HF vs LP-LF: p = <2e-16). The LP-LF treatment had significantly smaller shoals than all other treatments (Mann-Whitney U test: p < 0.001). The HP-LF and LP-HF treatment showed intermediate shoal sizes with the HP-LF treatment having a significantly higher mean shoal size than the LP-HF treatment (Mann-Whitney U test: p = 1.5e-06).

**Fig. 1.**
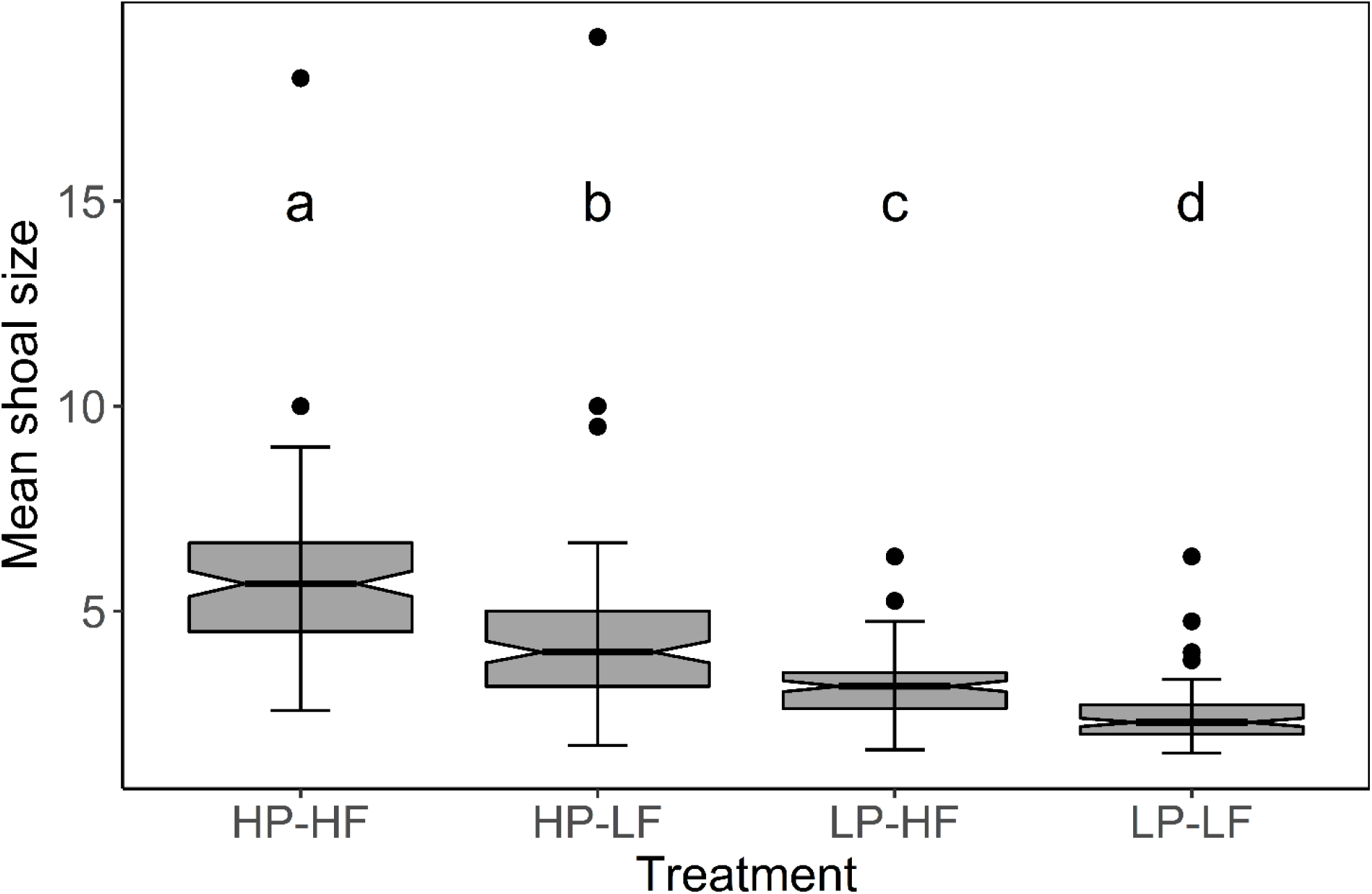
Mean shoal size across four treatments using the combined data of two sets of experiments. HP-HF represents high predation high food, HP-LF represents high predation low food, LP-HF represents low predation high food and LP-LF represents low predation low food. The thick horizontal line in the box depicts the median, and open circles represent outliers. The different letters above each box indicate significant differences (P < 0.001) between groups

A zero-inflated Poisson regression indicated that there were significant differences in the number of fish occupying different shoal size categories across treatments as opposed to a null model where there was no difference in the number of fish in each shoal size category across treatments (ΔAIC = −2402.284, p = <0.0001) (Fig 2). The HP-HF treatment had a higher proportion of fish in the largest shoal category (16–20 category) than in all other categories (vs 1–5: Estimate = 4.088013, z = 20.46354278, p = <0.05; versus 6–10: Estimate = 0.441803935, z= 5.540316392, p = <0.05; vs 11–15: Estimate = 0.066685531, z = 1.178214029, p = 0.099) (Table 1). The LP-LF treatment had a higher proportion of fish in the smallest category (1–5) than in the other categories (versus 6–10: Estimate = 4.977278, z = 20.98810376, p = <0.05; versus 11–15: Estimate = 5.666207042, z = 25.17957806, p = <0.05; versus 16--20: Estimate = 5.774575853, z = 25.89471241, p = <0.05) (Table 1).

**Fig. 2.**
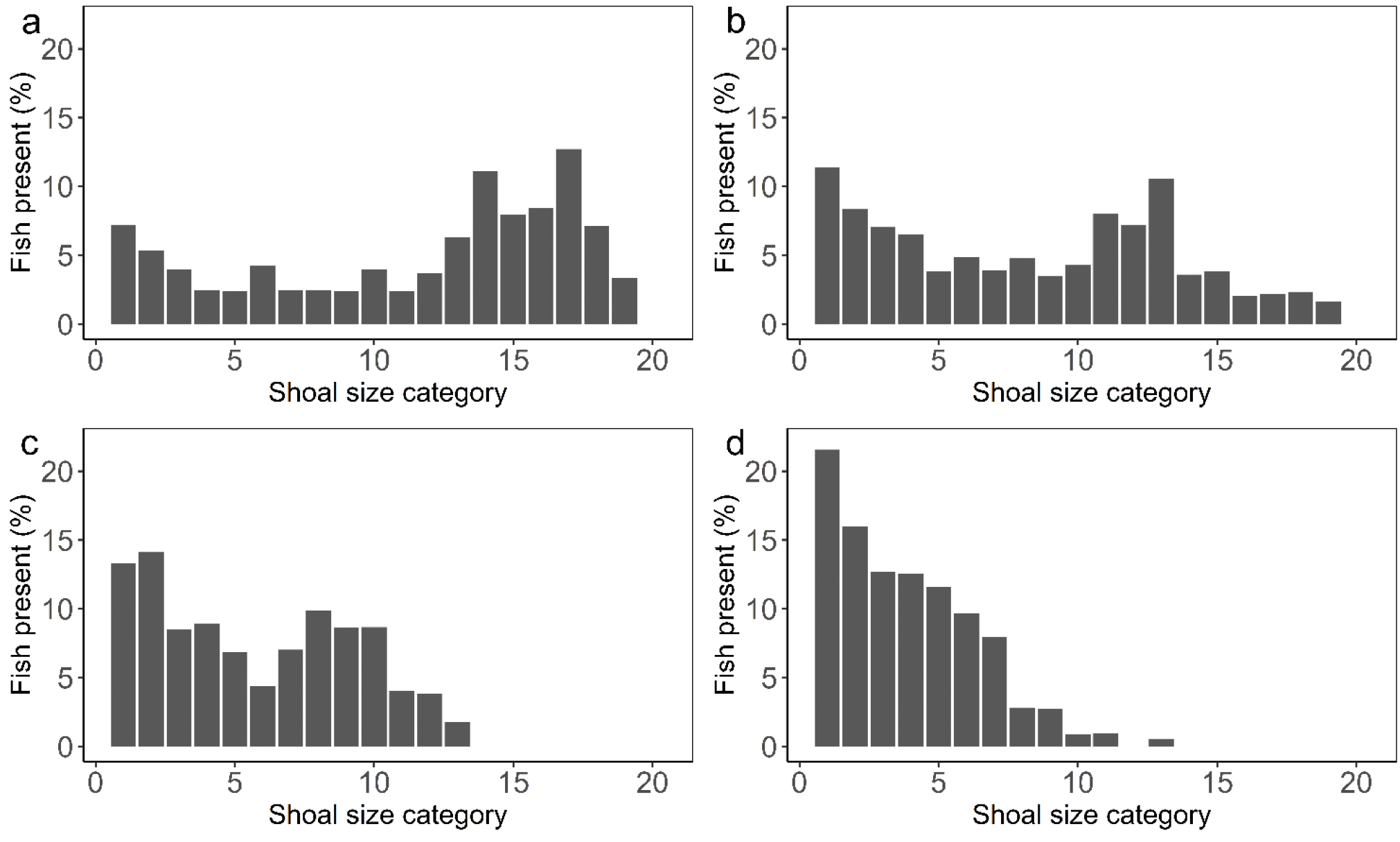
Percentages of fish occupying different shoal sizes under the four treatments A - high predation high food (HP-HF), B - high predation low food (HP-LF), C - low predation high food (LP-HF) and D - low predation low food (LP-LF)

**Table 1.** Results of the pair-wise comparison evaluating the frequency of fish occupying different shoal sizes (1-5, 6-10, 11-15 and 16-20) across the four treatments (High predation and high food availability: HP-HF; High predation and low food availability: HP-LF; Low predation and high food availability: LP-HF; Low predation and low food availability: LP-LF)

| Pair of groups | Estimate | Std. Error | z value | P value |
| --- | --- | --- | --- | --- |
| HP-LF 1-5 vs HP-HF 1-5 | 1.4135 | 0.2507 | 5.6388 | 2.03E-06 |
| HP-HF 16-20 vs HP-LF 16-20 | -0.0832 | 0.0578 | -1.4402 | 0.9884 |
| HP-HF 16-20 vs LP-HF 16-20 | 0.1129 | 0.0417 | 2.7096 | 0.3240 |
| HP-HF 16-20 vs LP-LF 16-20 | 0.1584 | 0.0364 | 4.3547 | 0.0014 |
| HP-HF 1-5 vs LP-LF 1-5 | -1.5282 | 0.2987 | -5.1156 | 3.64E-05 |
| HP-LF 1-5 vs LP-LF 1-5 | -2.9417 | 0.2716 | -10.8300 | 1.54E-13 |
| LP-HF 1-5 vs LP-LF 1-5 | -1.2471 | 0.2999 | -4.1576 | 0.0033 |
| HP-HF 1-5 vs HP-HF 16-20 | 4.0880 | 0.1998 | 20.4634 | 0 |
| HP-HF 16-20 vs HP-HF 6-10 | -0.4418 | 0.0797 | -5.5403 | 3.58E-06 |
| HP-HF 11-15 vs HP-HF 16-20 | 0.06668 | 0.0566 | 1.1782 | 0.9986 |
| LP-LF 1-5 vs LP-LF 6-10 | 4.9773 | 0.2371 | 20.9881 | 0 |
| LP-LF 1-5 vs LP-LF 11-15 | 5.6662 | 0.2250 | 25.1796 | 0 |
| LP-LF 1-5 vs LP-LF 16-20 | 5.7746 | 0.2230 | 25.8947 | 0 |

Shoal cohesion estimated using nearest neighbour distances indicated that the LP-HF treatment had significantly lower NNDs than that of other treatments (LP-HF vs HP-HF: Estimate = −0.100807, p = <0.05; LP-HF vs HP-LF: Estimate = - 0.092381, p = <0.05; LP-HF vs LP-LF: Estimate = −0.0848, p = 0.0001). However, NNDs did not differ significantly between other treatments (Table 2).

**Table 2.** Results of the comparisons of Nearest Neighbour Distances (NND) across the four treatments (High predation and high food availability: HP-HF; High predation and low food availability: HP-LF; Low predation and high food availability: LP-HF; Low predation and low food availability: LP-LF)

| Pair of groups | Estimate | Std. Error | z value | P value |
| --- | --- | --- | --- | --- |
| HP-HF vs HP-LF | -0.0084 | 0.0194 | -0.433 | 0.9727 |
| HP-HF vs LP-HF | -0.1008 | 0.0198 | -5.095 | < 1e-04 |
| HP-HF vs LP-LF | -0.0160 | 0.0195 | -0.817 | 0.8463 |
| HP-LF vs LP-HF | -0.0924 | 0.0196 | -4.705 | < 1e-04 |
| HP-LF vs LP-LF | -0.0075 | 0.0194 | -0.389 | 0.9800 |
| LP-HF vs LP-LF | 0.0848 | 0.0197 | 4.297 | 0.0001 |

## DISCUSSION

We find that the LP-HF treatment, which had low predation pressure and high food availability, had intermediate shoal sizes. On the other hand, fish in treatments that simulated high predation formed larger shoals both in low and high food conditions, while fish that experienced low predation and food (LP-LF treatment) formed smallest shoals (Fig. 1). Several studies have reported that fish tend to choose larger shoals when predator attacks are simulated (Magurran 1990; Magurran and Seghers 1991; Sogard and Olla 1997; Hoare et al. 2004; Huizinga et al. 2009). For example, Krause & Godin (1994) showed that banded killifish (*Fundulus diaphanus*) chose the larger of two possible shoals when an avian predator attack was simulated. Similarly, Magurran et al. (1987) reported that minnows (*Phoxinus phoxinus*) switched from dispersed small shoals to a single compact shoal when they detected predatory pike fish (*Esox lucius*). Larger group sizes have also been observed under high predation rates in other groups of animals such as mammals and birds (Fitzgibbon 1990; Ebensperger and Wallem 2002; Beauchamp 2004). The benefit of being part of a large group probably stems from the fact that attacks by predators are less successful in the case of larger prey groups (Landeau and Terborgh 1986).

Experiments have also shown that starved fish tend to choose smaller shoals compared to well-fed fish (Krause 1993; Reebs and Saulnier 1997; Hoare et al. 2004), suggesting that food deprivation causes fish to avoid competition incurred by being in a shoal (Hoare et al. 2004). Thus, shoaling behaviour can be maladaptive when foraging becomes a priority because of increased individual competition for food (Pitcher and Parrish 1993; Ryer and Olla 1995; Miller and Gerlai 2007). Overall, our study supports results from other studies wherein shoal size increased with predation and decreased with starvation (Magurran and Pitcher 1983; Magurran 1990; Hager and Helfman 1991; Dugatkin and Godin 1992; Krause 1993; Pitcher and Parrish 1993; Hoare et al. 2004).

Additionally, fish in the high predation and low food (HP-LF) treatment formed larger shoals than fish in the two low predation treatments (LP-LF and LP-HF) irrespective of their starvation level (Fig. 1 & 2). Although we cannot establish that the differences in selection pressure between the high and low predation conditions are the same as that of the starvation conditions, our results suggest that predation has a stronger effect on shoaling tendencies compared to food availability, at least under our experimental conditions. Previous studies have shown that guppies *(Poecilia reticulata)* from high predation habitats are less likely to exhibit exploratory movement for foraging than guppies from low predation habitats (Ioannou et al. 2017). Furthermore, foraging is reduced in minnows in the presence of a pike fish predator (Magurran 1990). Both of these studies further corroborate the idea that predation threat may have a stronger influence on shoaling tendency compared to starvation.

Although the LP-HF treatment had both small and intermediate shoal sizes, fish in this treatment had higher shoal cohesiveness than those in all other treatments irrespective of starvation or predation pressure. Although we expected the two high predation treatments (HP-HF and HP-LF) to have greater shoal compactness than the LP-LF group, we found no significant difference in shoal cohesion. Increased shoal size increases competition for other social factors such as mating (Alexander 1974; Hoogland and Sherman 1976) and also increases the risk of parasitism (Phillips 1924; Freeland 1976; Duffy 1983), which could lead to reduced shoal cohesiveness (Alexander 1974; Mikheev 2009; Hockley et al. 2014). Thus, the tendency of fish to stay close to each other due to predation pressure may not be strong enough to overcome the tendency to repel each other within the shoal in order to reduce competition for other social factors (Mikheev 2009; Shelton et al. 2015). On the other hand, the LP-LF treatment showed significantly reduced shoal cohesion compared to the LP-HF treatment, which could be due to the competition for food (Hoare et al. 2004; Miller and Gerlai 2007). We note that the distance between members of the shoal can be oscillatory (Miller and Gerlai 2008). Thus, estimating shoal cohesion based on snapshots of shoaling behaviour, as in our study, may lack some information on the shoal compactness. However, we have measured shoal cohesion based on multiple snapshots obtained from each video recording session to account for the variation within treatments.

Interestingly, we find that patterns of shoaling behaviours are consistent with other studies even when shoaling was measured under conditions where predation and starvation stressors are relaxed, indicating that prior experience of fish can influence shoaling tendencies. Several studies have shown that prior experience can have a significant impact on social behaviour of fish (Engeszer et al. 2004; McRobert et al. 2010; Song et al. 2011; Heathcote et al. 2017; Ioannou et al. 2017; Wang et al. 2019). For instance, it has been shown that fish in groups that have experienced predation had higher activity compared to naive groups while the activity of predator experienced singletons decreased compared to naive ones (Wang et al. 2019). Similarly, fish from high-predation habitats show reduced tendency to initiate explorative movements and show stronger homogenous movements compared to fish from low-predation habitats (Ioannou et al. 2017). Further, striped parrotfish (*Scarus iseri*) that have previously encountered high predation from native predators shows better resistance against a novel non-native-predator than fish experienced low predation via heightened risk avoidance (Berchtold and Côté 2020). In Zebrafish, it has been shown that they generally tend to prefer shoals of individuals which are visually identical to them (Rosenthal and Ryan 2005; Snekser et al. 2010). However, rearing conditions and experience with dissimilar phenotypes can alter such preferences (Engeszer et al. 2004). Here, we add to these studies by showing that previous environmental conditions experienced by fish can influence shoaling behaviour.

Many studies on the group behaviour of fish have tested the independent effects of predation and food availability on shoaling in fish (Magurran 1990; Hager and Helfman 1991; Krause 1993), whereas both predation and starvation often operate simultaneously to different extents in natural communities (Miller and Gerlai 2011). Our experiment, on the other hand, examines how prolonged exposure to different levels of combined effects of both predation and starvation can influence shoaling behaviour. Further, many studies exploring the factors influencing shoal dynamics use choice experiments in which a test fish that has undergone predation or starvation treatments is given visual cues of two shoals differing in size (Krause 1993; Krause and Godin 1994; Reebs and Saulnier 1997). However, the test fish in such choice experiments can neither interact with other members of the shoal nor access olfactory or auditory information from other members in the shoal (Miller and Gerlai 2007). Also, the test fish in these choice experiments are physically separated from the shoal and thus measure only forced-choice preferences based only on visual cues and, therefore, their interactions with the shoal can be questioned (Miller and Gerlai 2011). In our study, there are no restrictions to the choices that fish can make and they can freely interact with each other, and thus can spontaneously form shoals. Although some studies have examined how shoaling behaviour changes in the presence of both predatory and food cues (Godin 1986; Magurran et al. 1987; Morgan 1988; Godin and Crossman 1994; Hoare et al. 2004), these studies measure shoaling at the time when fish are subjected to these cues. On the contrary, we measure shoaling behaviour after prolonged exposure to predation and starvation and in the absence of these cues thus accounting for the dynamic nature of shoaling behaviour.

## CONCLUSIONS

Overall, our experiments suggest that exposure to prolonged environmental conditions such as predation pressure and resource availability can influence shoaling behaviour in zebrafish even when these conditions don’t persist. Further, we show that predation tends to have a greater influence than starvation pressure on shoal size, but not on shoal compactness. Such patterns could be because, compared to the failure to obtain food and mating, the failure to avoid predation can be more catastrophic to future fitness (Lima and Dill 1990). Thus, species should experience stronger selection pressure towards evolving anti-predatory strategies that reduce predation as compared to other activities. Our study provides insights on how prolonged exposure to environmental conditions can shape social behaviours and group dynamics.

## ACKNOWLEDGEMENTS

We thank Hema Somanathan for suggestions on the experimental design. We thank Rahul G Kumar, National Bureau of Fish Genetic Resources (NBFGR) and Jayapriya C.S. for information on housing and maintaining Zebrafish populations. We thank Anna Maria Dominic, Dilshad V.P. and Gopal Murali for helping with maintaining the fish populations. We thank Gopal Murali, Harshad Mayekar and Amal S. for discussions on the experimental setup. We also thank Noam Miller for discussions and suggestions.

## DECLARATIONS

### Authors Contributions

NAM and VPC conceived the study and designed the experiment; NAM carried out the experiment and analysed the data with inputs from VPC and UK; VPC prepared the figures; UK provided materials; NAM, VPC and UK wrote the paper and gave final approval.

### Ethical Approval

All applicable international, national and/or institutional guidelines for the care and use of animals were followed.

### Consent to Participate

All co-authors of the article are aware of their contributions and have agreed to being so named.

### Consent for publication

The authors guarantee that the contribution to the work has not been previously published elsewhere. All co-authors are aware of the fact and agreed to being so named.

